# PhysiCell training apps: A case study for creating interactive training materials for scientific software packages

**DOI:** 10.1101/2022.06.24.497566

**Authors:** Aneequa Sundus, Furkan Kurtoglu, Kali Konstantinopoulos, Mary Chen, Drew Willis, Randy Heiland, Paul Macklin

**Author notes:** These authors contributed equally to this work (Aneequa Sundus, Furkan Kurtoglu).

## Abstract

Cell-based tissue simulations require not only the ability to write new code in a simulation framework, but also an understanding of underlying mathematical models, background biology, and parameters for each behavior of an agent. This can entail a steep learning curve for interdisciplinary researchers joining computational biology research. We have created a suite of cloud-hosted open-source tools to separately explore and learn key components of an agent-based cellular simulation framework. This creates an self-contained environment to learn and test functions of cells and the micro-environment in a modular fashion before creating more detailed, research-focused simulation models.

## 1 Introduction

Multicellular systems biology is an interdisciplinary field. It requires knowledge of basic biology, physics, and mathematical modeling to transform domain knowledge into a mathematical format. To facilitate this transformation, the open source scientific community has developed modeling frameworks (e.g. Chaste[1], CompuCell3D[2], BioDynaMo[3], and HAL[4]) in a variety of programming languages. Once these frameworks’ generalized code bases are developed, specific models can be efficiently developed–often without understanding all the knowledge and code behind it. Many system modeling researchers join the field from technically diverse backgrounds, ranging from biology and informatics to engineering and mathematics. While simulation software packages are essential tools for working work in systems biology and computational biology, they may present a steep learning curve for scientists entering the field from diverse backgrounds. There are ongoing academic discussions seeking better ways to train researchers in computational modeling field. Madamanchi et al. [5] focused on challenges for natural science students as they venture into the computational modeling world. They identified that most such students seek to use computational modeling to answer domain-specific research questions that cannot be answered in wet labs. It shows that many scientists with the minimum computational background are potential users of computational modeling and simulation tools.Madamanchi et al. [6] discussed the need for user-friendly tools in computational modeling training, along with the potential of using focused web applications (“microapps”) for education and training. In this report, we demonstrate a suite of microapps to address the training needs of new multidisciplinary users of biological simulation software.

Agent-based simulation software (e.g.,PhysiCell[7], Chaste[1], and

CompuCell3D[2]) is widely used for creating cell based simulations. (See Metzcar et al. [8] for a review of cell-based models.) These models can be more intuitive to understand than differential equation-based models due to rule-based nature of assigning interpretable biological behaviors to cell agents. Each cell of same cell type behaves independently even though they share the same point with same rules and initial conditions; due to stochasticity (both biological and introduced in code), they soon desynchronize and follow individual trajectories. Cell-based models generally simulate the tissue (micro)environment for these agents/cells by integrating diffusion solvers in the background, thus modeling the secretion, movement, and consumption of chemical substrates. Thus, agent-based modeling frameworks provide rich, built-in capabilities, but can require extensive training and exploration to master.

## 2 Statement of Need

Many complex simulation software packages used in academia are open source and are managed by small research teams. For such small teams, training new people outside their labs can become a bottleneck to dissemination and adoption. One common solution is to arrange workshops to train new people, such as PhysiCell’s 2021 Virtual Woprkshop and Hackathon [9], recent CompuCell3D workshops [10], and the overall *Software Carpentry* approach [11]. There have been many efforts to create interactive materials for systems biology simulations. One earlier example of such training materials is *bugbox* [12]; this desktop-based predator-prey simulation application must be downloaded and installed before students can explore model parameters. In recent years, there have been efforts to create training materials more tailored to specific simulation software packages. In recent BrainIAK tutorials [13], the authors created learning materials for functional MRI analysis. They created Jupyter notebooks for users to run on their own to learn different concepts with open source data. They also created training materials around these notebooks to explain the concepts in fuller detail.

Although R is not frequently used to build simulation models, the Shiny package [14] can be used to transform R-based models into web applications for user-friendly sharing. For example, Handel [15] used R/Shiny to create an online, interactive systems immunology model. Their small, lightweight app allows users to explore different parameters in a model of infection and immune response. In “A virtual lab for teaching physiology” [16], David Granjon created an R/shiny app to dynamically explore the effects of different genes on calcium and phosphate homeostasis in rats and humans.

Generally, there are fewer such platforms for transforming Python-, JavaScript-, and C++-based computational models to shareable web applications. Artistoo [17] is a framework based on JavaScript that lets users build interactive, explorable simulation models of cells and tissues in a web browser, using a cellular Potts model [18] to simulate cells. Heiland et al. [19] developed *xml2jupyter* as a general framework to automatically create Jupypter-based graphical user interfaces (GUIs) for PhysiCell models built in C++, which can subsequently be shared as cloud-based web models on the nanoHub platform [20]. Similarly, CompuCell3D was recently adapted to hosting on nanoHUB [21]. Most of these applications provide specific examples for users to explore as a learning project. QUBES [22] also hosts a collection of training materials for qualitative biology. NetLogo[23] is another example for exploring mathematical models online by adjusting different parameters; however, NetLogo is not specifically tailored to biology.

Most of these applications do not explicitly link theoretical biological knowledge with programmable application. Instead, they generally assume that the user is already knowledgeable in those concepts or frameworks. Moreover, most of these frameworks are not physics-based (and instead restrict cells to non-realistic lattice positions with uniform size), while the cellular Potts-based models (e.g., CompuCell3D and Artistoo) are framed around interpreting biological behaviors as energy terms, which can be non-intuitive. There are unmet needs for online simulation frameworks where the cell behaviors are grounded in more realistic physics, cell behaviors can readily be associated with biologically meaningful parameters, and the models (and the underlying biology) can be taught through online interactive models that do not require software installation and additional coding.

## 3 Training Applications

We filled these unmet needs in training materials by creating a suite of small standalone applications that each focus on a different feature of cell-based models. A critical feature of our cloud-hosted interactive applications is that they are accompanied by training material to help the user understand the biological concept that each app presents. We have also designed our applications such that all parameters unrelated to a particular concept are fixed. This divide-and-conquer approach helps the user focus on the function of one parameter at a time. This way, we have divided the basic modeling framework into seven applications. Students will train on each aspect in each app and then see how all of these concepts work in relation to each other in a final simulation. This effectively provides scaffolding for new learners as they gain familiarity with the biological and modeling underpinnings.

The suite serves both to introduce key biological concepts of cell-based modeling while also specifically training users to run PhysiCell [7]: an open-source physics-based cell simulation framework.PhysiCell provides key cell functions (particularly cell cycling, death, mechanics, migration, secretion/uptake, as well as newer immune functions) that can be used to write complex simulations of cells in their environment. PhysiCell provides support to run these simulations on desktops as well as on clusters and high performance computing (HPC) resources. The core code of Physicell is written in C++, with input files provided in extensible markup language (XML), and output files stored as XML and MATLAB files. Beyond the core framework is a growing software ecosystem, including packages like xml2jupyter [19] that that can help run the compiled models in a Python-based Jupyter Notebook, PhysiBoSS [cite] (integrated Boolean signal networks), PhysiCool [cite] (), and PhysiCell [cite] (HPC extension of PhysiCell). With its wide array of programming languages, file formats, and packages, PhysiCell can present a steep learning curve not only for students new to systems biology, but also those who are experienced with systems biology but unfamiliar with this specific software.

Our training applications are hosted and available without charge on NanoHUB: an online platform to host simulations and other software tools [20]We selected seven applications because they capture the core functionality of PhysiCell. While these introductory applications do not completely capture the breadth of functionality offered by PhysiCell, they are sufficient to train a user to create complex multi-cellular simulations. Moreover, after learning these key core behaviors, learners are well-equipped to learn new features.

Each training app has an graphical user interface (GUI), as depicted in Figure In the About tab, we describe the biological and mathematical background of the app, along with some suggested parameter sets for exploration. This is followed by the `Config Basics, Microenvironment`, and `User Parameters` tabs. These three tabs provide different customization options to users, including parameters that can be modified to explore in the app’s simulation model. We have minimized the number of changeable parameters to help focus the learner’s attention on the selected modeling aspect for each app. Details about the theory and application of these applications are described below.

### 3.1 Chemical Diffusion

Extracellular diffusion and decay are critical to cell-cell communication (through secreted chemical factors) [cite 10.1007/978-3-319-42023-3 12], as well as drug transport in tissues. Therefore, it is critical that modelers explore and understand the relationships between diffusion, decay, secretion, uptake, boundary conditions, and penetration distance. PhysiCell [7] uses BioFVM [24] to simulate the chemical environment of cells. BioFVM [24] is a diffusion transport solver that allows substrate diffusion models to be run on modern desktop computers as well as supercomputers. In this app, we demonstrate the chemical environment using two substrates: oxygen and ChemicalA. Users can modify the diffusion and decay rates of each substrate to obtain different diffusion length scales 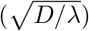. Users can also set the boundary conditions for each substrate conditions to either von Neumann (zero flux) boundary conditions or Dirichlet (fixed value) boundary conditions. Within the Dirichlet boundary parameters, there is an option to fine-tune the values for individual boundaries. Users can also add one Dirichlet node (a location where the substrate value is held fixed, similarly to a boundary condition) anywhere in the domain. Figure 2a shows the distribution of oxygen in the microenvironment after 6 hours and 30 minutes of simulated time, with Dirichlet conditions of 100 on the positive x-axis and 0 on the negative x-axis, and zero flux (von Neuman) conditions on the remaining boundaries. There is also a Dirichlet node present in the domain at (100,100) with a concentration of 100 on it. No chemical is being absorbed by any cell in the environment in this demonstration app, allowing users to focus solely on the impact of diffusion, decay, and boundary conditions. The Chemical Diffusion app is accessible at https://nanohub.org/resources/microenvnmtr.

**Fig. 1:**
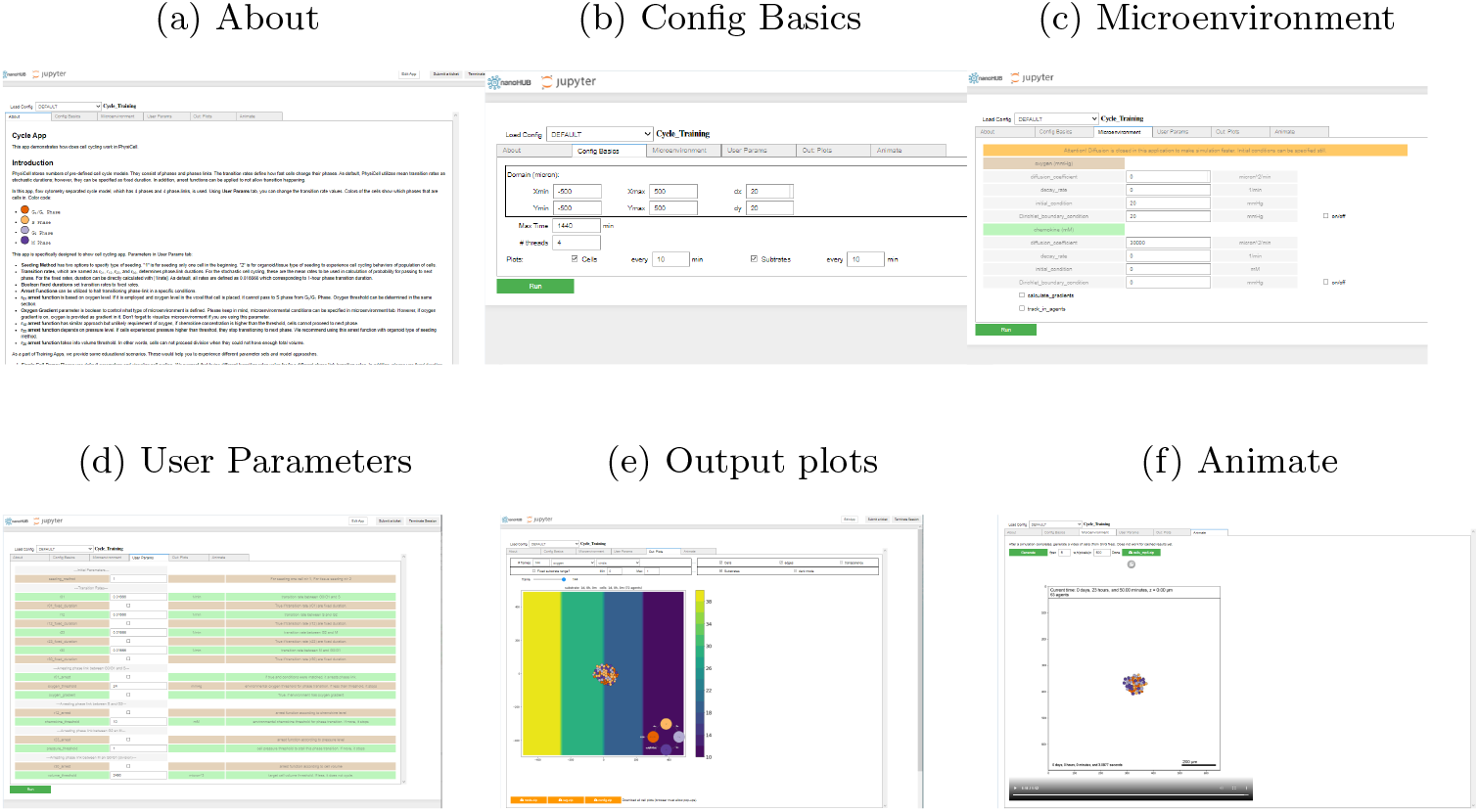
User Interface of each Training App. **(a)** The About tab includes an introduction to the app and some suggested scenarios for the user to get familiar with app and its educational concepts. **(b)** The Config Basics tab is where the user can set the domain size of simulation and how often data is being stored and displayed. **(c)** The `Microenvironment` tab displays chemical substances present in the environment along with their diffusion and decay rates. Users can also set different boundary conditions for these substrates. **(d)** The `User Parameters` tab is the main focus point for most applications. Users can set parameters related to the module under discussion and then run the model by clicking on “Run” button. **(e)** In `Plots`, the user can observe the behavior of cells and substrate diffusion in the environment over time. **(f)** In `Animate` tab, users can make small animation of all outputs for rapid dynamic visualization.

**Fig. 2:**
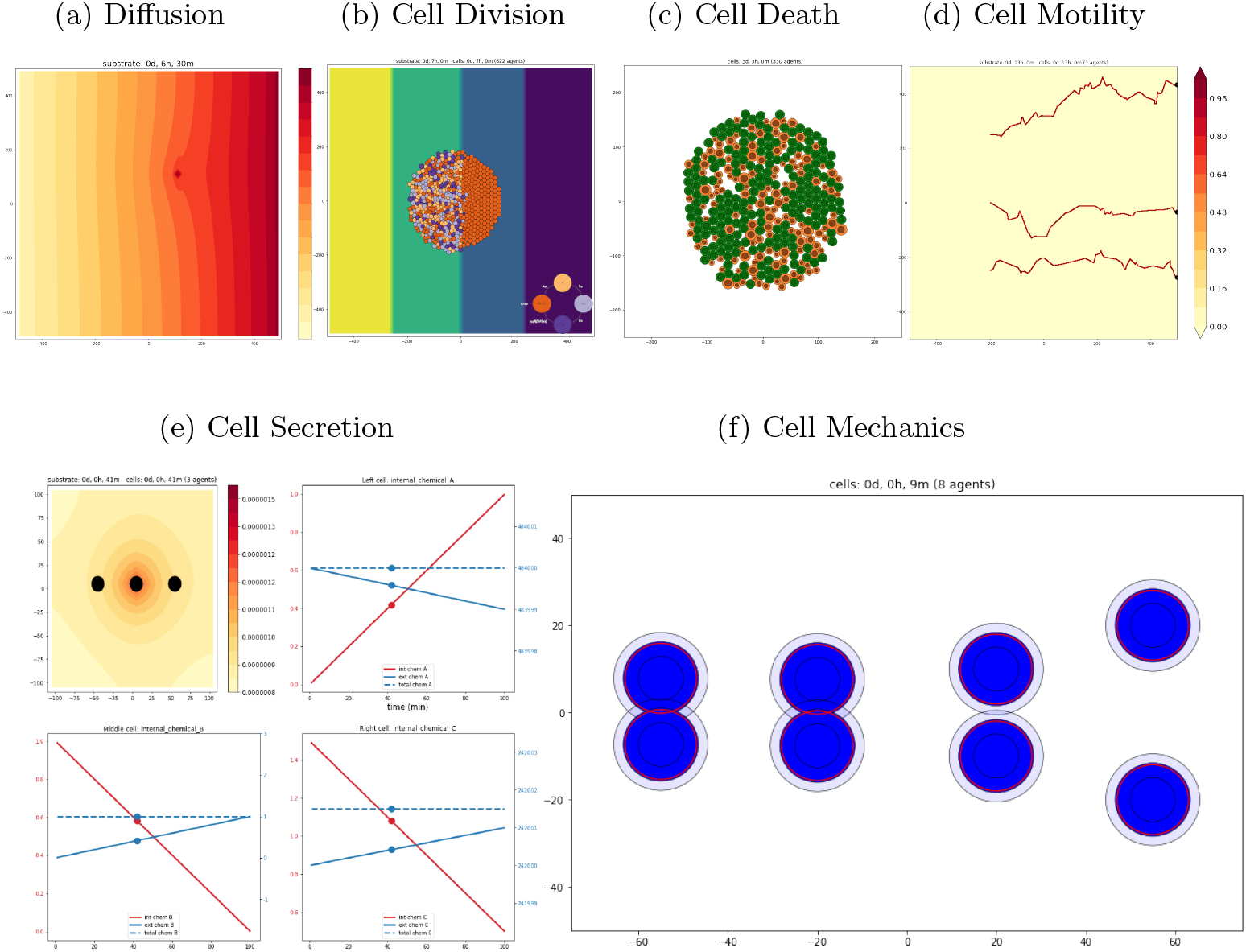
Typical Outputs for the Applications. **(a)** Chemical Diffusion shows bands of chemical gradient formed by introducing Dirichlet boundary Conditions on the x-axis. **(b)** Cell Division displays cells going through different color coded-cell cycle phases according to their microenvironment **(c)** Cell Death shows an apoptosis cell death model introduced on a blob of cells **(d)** Cell Motility shows the biased random path of three motile cells **(e)** Cell Secretion displays three cells with their relative chemical secretion and absorption. Graphs display the amount of chemicals inside and outside the cell for each one **(f)** Cell Mechanics shows four pairs of cells placed at different distances resulting in different adhesive and repulsive forces for each pair.

### 3.2 Cell Division

Because cell division is crucial to cancer biology, developmental biology, and selective pressures in multicellular ecosystems, modelers should understand the structure and behavior of common cell cycle models, and how they manifest in cell-based models. Cell division is modeled in PhysiCell through different cell cycle models. Each cell cycle model consists of several phases (e.g., G_0_/G_1_, S, G_2_, M), and cells transition from one phase to the next based on user-provided transition rates. These rates can be coupled with some external factors to control the cell cycle. PhysiCell also provides some arrest functions that can lock the cell in a particular phase. At the end of the final stage, the cell divides into two daughter cells. In this app, we use PhysiCell’s built-in `flow cytometry separated` cycle model (where cells transition from *G*_0_*/G*_1_ to *S* to *G*_2_ and finally to *M* phase) to explore cell division. In the app, four factors control the four transition rates between cell cycle phases in the model. Users can control target concentrations of substrates, volume, and pressure to control transition from one phase to the next. Figure 2b shows the output of the cell cycle app, where cells are prevented from transitioning from *G*_0_*/G*_1_ to *S* if the local oxygen value is less than 25 mmHg. We can see that cells in the left half of the domain with oxygen concentration higher than 25 mmHg continue to progress through the cell cycle phases and dividing, while cells on the right are arrested in *G*_0_*/G*_1_. Also notice that the cells present on left side are of the same type, yet they are not synchronously progressing through the cycle transitions. This is due to stochastic behavior introduced in the code to observe more realistic behavior. It is important to note that diffusion is not enabled for this app, so substrate gradients are not responsible for the observed variation in cell behaviors. The Cell Division app is accessible at https://nanohub.org/resources/trcycle.

### 3.3 Cell Death

Cells can undergo a broad range of types of death, including apoptosis (controlled cell death, and generally not inflammatory), necrosis (the prototypical uncontrolled death due to injury or energy collapse, typically driving inflammatory responses), and other death processes as described in [25]. These can drive a broad range of changes in multicellular systems. PhysiCell includes built-in models of two of these forms of cell death: apoptosis and necrosis. Apoptosis is programmed cell death, which helps maintain regular tissue growth and function. In contrast, necrosis is un-programmed cell death due to injury or disease. The apoptosis model has only a single “apoptotic” phase, primarily characterized by cell shrinkage. The necrosis model has 2 phases: the swelling phase and the lysed phase. The cell enters the swelling phase and grows in size until it reaches a specific “bursting”volume, after which it lyses (bursts) and slowly shrinks (and potentially calcifies). Figure 2c shows cell death with the apoptosis model. Brown cells are going through apoptosis. They swell before rupture. Green cells have not yet undergone apoptosis. This also shows the stochastic behavior in the model, since cells do not go through phases of apoptosis all at once. We also provide option to simulate necrosis death in Cell Death training application. Users can control death rate and biomass change rate to visualize both death models. The Cell Death app is accessible at https://nanohub.org/resources/trdeath.

### 3.4 Cell Motility

As opposed to motion under the influence of adhesive and repulsive forces, cell motility referes to active cell locomotion or migration, either randomly or in response to some external stimulus (e.g., via chemotaxis). In PhysiCell (as well as many other cell-based models), motility is modeled via a biased random walk: a cell chooses a direction (either along a migration bias direction, randomly, or a combination based on its bias), move along that direction for a duration governed by its migration persistence time, and then choose a new direction of travel. Because cell motility is a key driving factor in cancer invasion, tissue morphogenesis, and immune cell interactions, it is crucial that learners can model and explore these functions. Biased random migration–and its parameters–can be difficult and abstract to understand, thus motivating a need for interactive training and exploration. In this app, users can control cell motility by setting the migration bias, migration speed, migration bias angle, and persistence time. We display each cell’s path in this training app using a trail of a non-diffusing chemical “tracer” in the microenvironment. At the end of the simulation, the effect of different parameters on cell movement can be observed by studying the cell paths. Figure 2d shows the path of three cells in the domain when they move toward a 30-degree x-axis gradient with a bias of 0.6. The Cell motility app is accessible at https://nanohub.org/resources/trmotility.

### 3.5 Cell Secretion And Uptake

Cell secretion and uptake of diffusable substrates mesh with diffusion and decay (see above) in driving intracellular chemical communication. Thus, multicellular biologists stand to benefit from exploring the parameters driving these key processes and their impact on extracellular substrate concentration fields, as well as changes in intracellular substrate values. This app demonstrates the secretion and uptake functionality of cells in the PhysiCell environment. Cells remove (or consume or uptake) chemicals from the microenvironment based on their uptake rates, or secrete them. PhysiCell tracks the total amount of intracellular substrates in each cell; conservation of mass dictates that any secreted substrate is balanced by a corresponding reduction in intra-cellular substrate, whereas and uptake from the microenvironment is balanced by a corresponding increase in the intracellular substrate. Taken together, mass conservation is guaranteed to keep the total sum of chemical within the cell and in the microenvironment constant. It is challenging to visually show the amount of chemicals in cells and outside due to difference in scales of the amount present inside and outside of cells. We have solved this difficulty by separately plotting the amount of chemicals inside cells and outside in the environment. Figure 2e shows a sample output of the application. There are three cells in the simulation. The left cell uptakes Chemical A from the environment, the middle cell secretes Chemical B, and the right cell secretes Chemical C. Three graphs show the amount of chemicals present in each cell and the microenvironment. We can see that the total amount of chemicals remains constant due to mass conservation. Parameter values used for this output can be seen in scenario 2 of the app About page. The cell Secretion application is accessible at https://nanohub.org/resources/trsecretion.

### 3.6 Cell Mechanics

A defining feature of cell-based models is that they track the movement of individual cells, often by tracking the net balance of adhesive and “repulsive” forces acting upon each cell to determine their resulting velocities [cite Metzcar et al review]. Multicellular biologists who are switching from cell population kinetics to spatial modeling for the first time should therefore explore how individual cell mechanical parameters contribute to overall cell movement. Cell mechanics in Physicell is modeled by simulating adhesive and repulsive forces (which model mechanical resistance to deformation), similarly to to the Leonard-Jones potential in atomic and particle physics.

The range and strength of these forces ultimately create an equilibrium position for a cell at a certain distance from its neighbors, at which attractive adhesive forces between the two cells balance with repulsive forces to yield a net zero displacement velocity. In the training app, we created four pair of cells seeded at different distances from each other. Users can modify the adhesive and repulsive forces as well as the maximum interaction distance of these forces to explore their impact of the emergent cell velocities and positions. The interaction radius of each cell is displayed using a blue field around it, while the (parameter-dependent) equilibrium distance is denoted by red circles. Any cell pairs that are within their interaction radii (i.e., their blue circles overlap) will pull each other towards the equilibrium spacing, while any cell pair beyond that interaction distance will not. When two cells have reached equilibrium, their red circles will just touch but not overlap. Notice that if any two cells are closer than their equilibrium spacing, they will push each other apart. After just 9 minutes of simulation, two right pair of cells are pushing each other; see Figure 2f. The third pair from right is getting attracted, while the leftmost pair is not moving since both are beyond the interaction distance. Cell Mechanics application is accessible at https://nanohub.org/resources/trmechanics.

### 3.7 Cell Volume

Cell volume in PhysiCell is used to capture liquid and solid fractions in the cytoplasm and cell nucleus. The cell will grow or shrink towards its steady-state (or “target”) volume, based on its individual stored parameters; its current volume is stored as a state. This also includes parameters to control biomass change rate in both nucleus and cytoplasm. Each cell has a target value for these parameters, which can be achieved at a given biomass change rate. Within this application, we have divided the data members of cell volume into two main categories: Parameters and State Variables. Parameters of the cell are those data members that define the steady-state volumes and biomass change rates. State Variables are used to store transitory values for the cell over time. In the time course, the cell will always try to return to its steady state. In this application, we have used a red circle to denote the cell’s steady-state size as determined by its parameters. Cells are color-coded according to a fluid fraction of cell cytoplasm and cell nucleus. Users can perturb the cell’s state and watch it evolve back towards its steady state, or adjust parameters to change the steady state and thus direct the dynamics. Even if you change the state variables and set them to inconsistent values, the cell will reach its steady state sooner or later. Cell volume represents a single cell and its nucleus. It is color-coded according to the fluid fraction of the cell cytoplasm and cell nucleus. Its behavior is more dynamic, showing how the cell goes to its target values from wherever its state variables are. The Cell Volume application is accessible at https://nanohub.org/resources/volumetr.

## 4 Results

Our goal for this work was to create user-friendly modules to train novice researchers on a cell-based modeling framework. We created this suite of seven open-source and made them broadly accessible with cloud hosting. Each app can be run independently, allowing users to select and focus on a single topic. We endeavored to make these applications faster by removing any functionality unrelated to the topic under discussion. Thus, these applications run fast, and we can see results almost instantly. We also provide a guide for the exploration on the front page of all applications, directing users to explore and outlining what they can expect from changing different parameters.

During the development of these applications, we worked in close collaboration with the principal developer of the software and the principal developer of user a interface for cloud-hosted applications. This close in-house collaboration resulted in improving not only training applications but also the underlying software. We identified the need in the software to quickly set Dirichlet conditions of each boundary, which was then implemented and added to the next release of PhysiCell. During the development and testing of these applications, we found and reported bugs in PhysiCell’s implementation of older function definitions, thus showing the integral role that the development of training software can play in quality control. We also worked with the developers of the Jupyter interface, which led to the addition of extra usability-focused features, such as adding separation tabs to separate different sets of parameters, adding an animate tab, and showing cell paths. We shared our initial versions of applications in our weekly lab meetings. Feedback from other lab members led us to explore some features in color template selection for applications. It is essential for applications like the cell cycle, where around half a dozen colors represent different stages in the cell cycle. Feed-back from lab members led us to refine the usability and explore colorblind-friendly color schemes. We decided to go with color palettes described in [26] for these applications. This showed the power of integrating the development of training applications and user interfaces directly with day-to-day lab operations.

Our initial development of these applications began as a side project with the aim of creating some basic training materials for the first-ever PhysiCell workshop. However, the scope and scale of the project soon expanded, providing us with an opportunity to learn how to decouple a complex software system into manageable pieces, as well as how to select appropriate parameter sets and models to represent those components. Additionally, we incorporated customized visualizations into each application to highlight their specific points. This endeavor required significantly more time and resources than we had initially anticipated, which raises the issue of the need for appropriate funding resources. As we have previously mentioned, training materials are an essential component of the scientific software ecosystem, and it is imperative to provide adequate funding avenues to develop such tools and training materials, rather than treating them as an afterthought.

## 5 Discussion

We used these applications as teaching aids in an introductory class to systems biology modeling, and to introduce agent-based modeling to undergraduate researchers. These applications helped in translating theoretical concepts into tools that could drive hands-on exploration by students. These applications were also included as teaching materials for the PhysiCell workshop [9], and they helped accelerate the learning curve for participants in the one-week program. These applications were heavily used in training sessions about microenvironment setup and cell phenotype. All the participants of the workshop were able to create their own PhysiCell models by the end of the workshop. These applications can also be used in multiple settings, such as high school-level biology classes to introduce concepts like diffusion, chemotaxis, and mechanical forces. We note that such applications could present an additional resource to include with documentation of other scientific software user guides, particularly for interdisciplinary fields where ideas from different disciplines intersect in one model.

Although these applications encompass the basic modules of a small cellular simulation, they cannot fully capture some of the more advanced features in PhysiCell that are used to write complex multi-scale simulations. While it is difficult to write narrowly tailored training applications for these features, intermediate-level users (after having learned the essential features in this training suite) can continue learning from numerous intermediate-level and advanced projects available on nanoHUB. We recommend using cancer bio-robots [27] and PhysiCell invader-scout-attacker system simulation [28] on nanohub for intermediate-level exploration of capabilities of PhysiCell. Three-Type Multicellular Simulation Lab [29] shows a very detailed and customized simulation in which almost all the parts are available to customize. Thus, this provides an advanced-level application for exploration. applications presented in this paper will serve as fundamental building blocks to move towards more complex simulations using PhysiCell platform.

The principle of accessibility is a fundamental principle in MathCancer lab. Our efforts have been concentrated on ensuring cross-platform and backward compatibility, as well as ever-evolving user interface design, to facilitate ease of use for a diverse community of PhysiCell users. These efforts have resulted in the creation of PhysiCell Studio[30], as well as the recent addition of a rule-based modeling language in Physi-Cell release 1.11.0 [31]. PhysiCell Studio is a graphical user interface (GUI) application to create and edit a PhysiCell model, run a simulation, and visualize results. The rule-based modeling language is a simple API that facilitates the building of models using cell signals and cell behaviors without any programming requirements. These developments raise a valid concern regarding the potential benefits of creating such applications for user training. Our answer is affirmative, as the inclusion of user interfaces on top of the core code introduces a level of abstraction. As users become more separated from the core library, it becomes increasingly important to provide them with the tools to comprehend the inner workings of the code, and to make informed modeling decisions. While an improved user interface may facilitate more efficient model implementation, the most critical aspect of modeling is always the design of the model. Without adequate training in the underlying assumptions and workings of the code base, mistakes are prone to arise. Therefore, the importance of creating standalone applications that are specifically designed to teach the inner workings of the code base cannot be overstated.

It should also be noted that training applications can be tailored to educational needs, with features such as focused views to block out distractions and prevent learner overwhelm. They provide a designed scaffolding that includes the first investigation by using default parameters followed by suggested exploration that is provided with each application. We also provide tailored visualization that is not general-purpose, but specific to the lessons of the training application. These specialized applications can serve distinct and complementary pedagogical goals, above and beyond the general aim of making the software accessible and easy to use.

## 6 Future Work

PhysiCell is constantly evolving with the addition of new features, such as advanced chemo-taxis, spring attachments, cell-cell contact, signals, and behaviors, since the development of these applications. This necessitates the creation of new training applications to introduce these concepts. With the release of PhysiCell version 1.10.4, new and complex functions were introduced, such as immune and interaction functions, phagocytosis, fusion, and transformation. These functions are highly intricate and require tailored training programs to comprehend. For example, modeling immune response involves defining effector attack and damage response, as well as vectors of attack rates and immunogenicity. Another recently added API, signal-behavior-response, can also benefit from customized training applications

As integrated biology becomes increasingly complex, the need for focused training materials becomes more apparent and important. The usability and training of the software drive its use, which in turn leads to a greater demand for more features. Nevertheless, the incorporation of more features requires a concurrent improvement in usability and training.

## Acknowledgments

We want to acknowledge and thank fellow Ph.D. students; John Metzcar, Yafei Wang, post docs; Heber Rocha and Michael Getz: and past and present undergrad members of MathCancer lab for their feedback and suggestion during the development of these applications. Their feedback on earlier versions of these applications was key in improving these applications and shaping them into their present form.

## Declarations

### Funding

We thank the Breast Cancer Research Foundation, Jayne Koskinas Ted Giovanis Foundation for Health and Policy, National Science Foundation (1818187 and 1720625), and National Cancer Institute (1U01CA232137-01) for generous support.

### Conflict of interest

The authors declare that they have no competing interests.

### Authors’ contributions

AS and FK led planning, development and refinement of the apps. KK, MC, and DW contributed to app development. RH provided technical support and refinement for Jupyter interfaces. AS, FK, and RH co-mentored undergraduate researchers. PM conceptualized and directed the work, obtained funding, and provided technical support for PhysiCell. AS led manuscript preparation, and all contributed to manuscript revision.

## Notes

### Competing Interest Statement

The authors have declared no competing interest.

### Summary of Updates

Added more discussion and detail in the effectiveness of training materials.

